# Global environmental drivers of local abundance-mass scaling in soil animal communities

**DOI:** 10.1101/2022.06.29.498075

**Authors:** Ana Carolina Antunes, Benoit Gauzens, Ulrich Brose, Anton M. Potapov, Malte Jochum, Luca Santini, Nico Eisenhauer, Olga Ferlian, Simone Cesarz, Stefan Scheu, Myriam R. Hirt

## Abstract

The relationship between species’ body masses and densities is strongly conserved around a three-quarter power law when pooling data across communities. However, studies of local within-community relationships have revealed major deviations from this general pattern, which has profound implications for their stability and functioning. Despite multiple contributions of soil communities to people, there is limited knowledge on the drivers of body mass-abundance relationship in these communities. We compiled a dataset comprising 155 soil-animal communities across four countries (Canada, Germany, Indonesia, USA), all sampled using the same methodology. We tested if variation in local climatic and edaphic conditions drives differences in local body mass-abundance scaling relationships. We found substantial variation in the slopes of this power-law relationship across local communities. Structural equation modeling showed that soil temperature and water content have a positive and negative net effect, respectively, on soil communities. These effects are mediated by changes in local edaphic conditions (soil pH and carbon content) and the body-mass range of the communities. These results highlight ways in which alterations of soil climatic and edaphic conditions interactively impact the distribution of abundance, and thus energy, between populations of small and large animals. These quantitative mechanistic relationships facilitate our understanding of how global changes in environmental conditions, such as temperature and precipitation, will affect community-abundance distributions and thus the stability and functioning of soil-animal communities.

## Introduction

Global alterations in environmental conditions are expected to have severe impacts on species communities and their contribution to our society (IPBES, 2019). In particular, soil-animal communities have important functions in many of nature’s contributions to people (NCP), including the decomposition of dead organic material, the recycling of nutrients, carbon sequestration and pest control (Blankinship et al. 2011, Bardgett and Van Der Putten 2014, Pereira et al. 2018). Many of these contributions can be quantified using fluxes of energy and material through the food webs, which strongly depend on both the distribution of body masses (i.e. the weight of an individual) and abundances (number of individuals per unit area) across species (De Ruiter et al. 1995, Neutel et al. 2002, Barnes et al. 2016, 2018, Jochum et al. 2021a). Therefore, fluctuations in the community composition and species’ relative densities in soil communities affect the flux of energy through the trophic levels (Schwarz et al. 2017) and, consequently, trophic multifunctionality (Potapov et al. 2019). Despite increasing evidence of direct effects of global warming and altered precipitation on soil biota (Blankinship et al. 2011, Yin et al. 2020), we know little about how these changes in environmental conditions modify the distributions of body masses and abundances within communities, which have strong indirect effects on ecosystem stability and functioning (Winfree et al. 2015, Wang and Brose 2018, Potapov et al. 2019, 2021). This knowledge gap hampers our ability to predict future NCP of soil communities.

Body size is a fundamental trait that regulates species’ biological rates, such as metabolism, biomass production, and feeding, and thus ultimately abundances (Peters and Wassenberg 1983, Woodward et al. 2005, White et al. 2007a). Generally, the body mass-abundance relationship is very consistently described, with density (N) decreasing with population-level body mass (M) following a negative three-quarter power law (Damuth 1981, 1987, Enquist et al. 1981, Allen et al. 2002). Studies using global or cross-community datasets that aggregate body masses and abundances from different local communities have provided ample empirical support for this relationship (White et al. 2007b, Hatton et al. 2019). However, studies describing this body mass-abundance relationship in local communities found deviations from the general negative three-quarter power-law scaling (Cyr et al. 1997, Cohen et al. 2003, Reuman et al. 2009, Gjoni and Glazier 2020), possibly related to gradients of human impact (Munn et al. 2013, Santini and Isaac 2021). This variation implies that local factors can modify the globally stable distribution of abundances across the size classes of species and therefore change local community structures, energy flux, and NCP.

There is extensive evidence for the general importance of environmental conditions, such as soil temperature, carbon content, or litter stoichiometry, for soil-animal abundances at different spatial scales (Ehnes et al. 2014, Ott et al. 2014, Phillips et al. 2019, Johnston and Sibly 2020). As these studies lump data across individual communities to derive a single body mass-abundance scaling relationship, we still know little about how these factors drive the scaling relationships within local communities, as described, for example, by the slope of the relationship. Yet, this is critical to understand how changes in the environment affect the energy distribution in communities and thereby ultimately the provision of ecosystem functions. Few comparisons of body mass-abundance slopes among soil communities showed variation depending on land-use types, soil acidity and stoichiometry, and the communities’ range in body masses (Mulder and Elser 2009, Ulrich et al. 2015). Nevertheless, none of these studies teased apart the direct and indirect pathways of these impacts on local body abundance-mass relationships in general and the importance of climate variables in particular.

We addressed this topic by synthesizing the so-far largest dataset on abundances and body masses in 155 soil invertebrate communities across different continents (Canada, Germany, Indonesia, USA). All local communities were sampled using the same methodologies to assess meso- and macrofauna (soil invertebrates in the body size range from ca. 0.5 mm to ca. 5 cm), and the resulting body mass-abundance relationships are community-specific. We hypothesized that soil temperature and soil water content, which are strongly dependent on climatic factors, have a major impact on body mass-abundance relationships in local soil communities (Johnston and Sibly 2020). We expected that these environmental variables exert a direct effect on edaphic conditions, such as soil pH and soil carbon content (Onwuka 2018, Hartley et al. 2021), thereby indirectly affecting the slopes of the local body mass-abundance relationships. Additionally, we also expected indirect effects on the body abundance-mass slopes mediated via changes in the body-mass range realized in the local communities (Ulrich et al. 2015). Overall, our study thus aims at disentangling the direct and indirect pathways how climatic and edaphic conditions affect the local distribution of abundances across size classes of soil animals.

## Material and methods

### Study sites

We investigated forest soil invertebrate communities from four globally-distributed geographic locations covering diverse environmental conditions (Figure 1). We compiled data from three large-scale projects: (1) the Biodiversity Exploratories project is located in the south-west, center, and north-east of Germany (Fischer et al. 2010). A total of 48 plots were sampled between 2008 and 2011. The habitats comprise beech and coniferous forest sites, and different land-use types: from intensively managed coniferous monocultures to nearly natural beech forests (see Ott et al. 2014 for a detailed description). The mean percentage of water content in the soil is 28% (in wet weight) and the mean annual soil temperature is 6.6°C. (2) The ECOWORM project was conducted across four northern North American forests in Canada and the USA between 2016 and 2017 (Eisenhauer et al. 2019), and a total of 80 plots were sampled. The forests in Canada (Barrier Lake North, Barrier Lake South and Bull Creek Hills) are situated in the Canadian Rocky Mountains, Kananaskis Valley, and are dominated by aspen tree species. The mean soil water content and temperature are 31% and 5.4°C, respectively. The US forest is located in northern Minnesota and has a mean soil water content of 26%, and a mean soil temperature of 8.1°C. The sites are covered by mesic forests and are dominated by sugar maple species. (3) 30 research plots of the collaborative German-Indonesian research project CRC990/EFForTS were set up in Jambi province in 2013, Sumatra, Indonesia (Drescher et al. 2016); the sites cover different land-use systems, from rainforest to monoculture rubber and oil palm plantations. Mean soil water content was 40% and mean soil temperature 24.6°C. A detailed description of the sampling methods applied can be found in the original studies (Ott et al. 2014, for sites in Germany, Barnes et al. 2014 and Potapov et al. 2019, for sites in Indonesia, Jochum et al. 2021b, for sites in USA and Canada).

**Figure 1:**
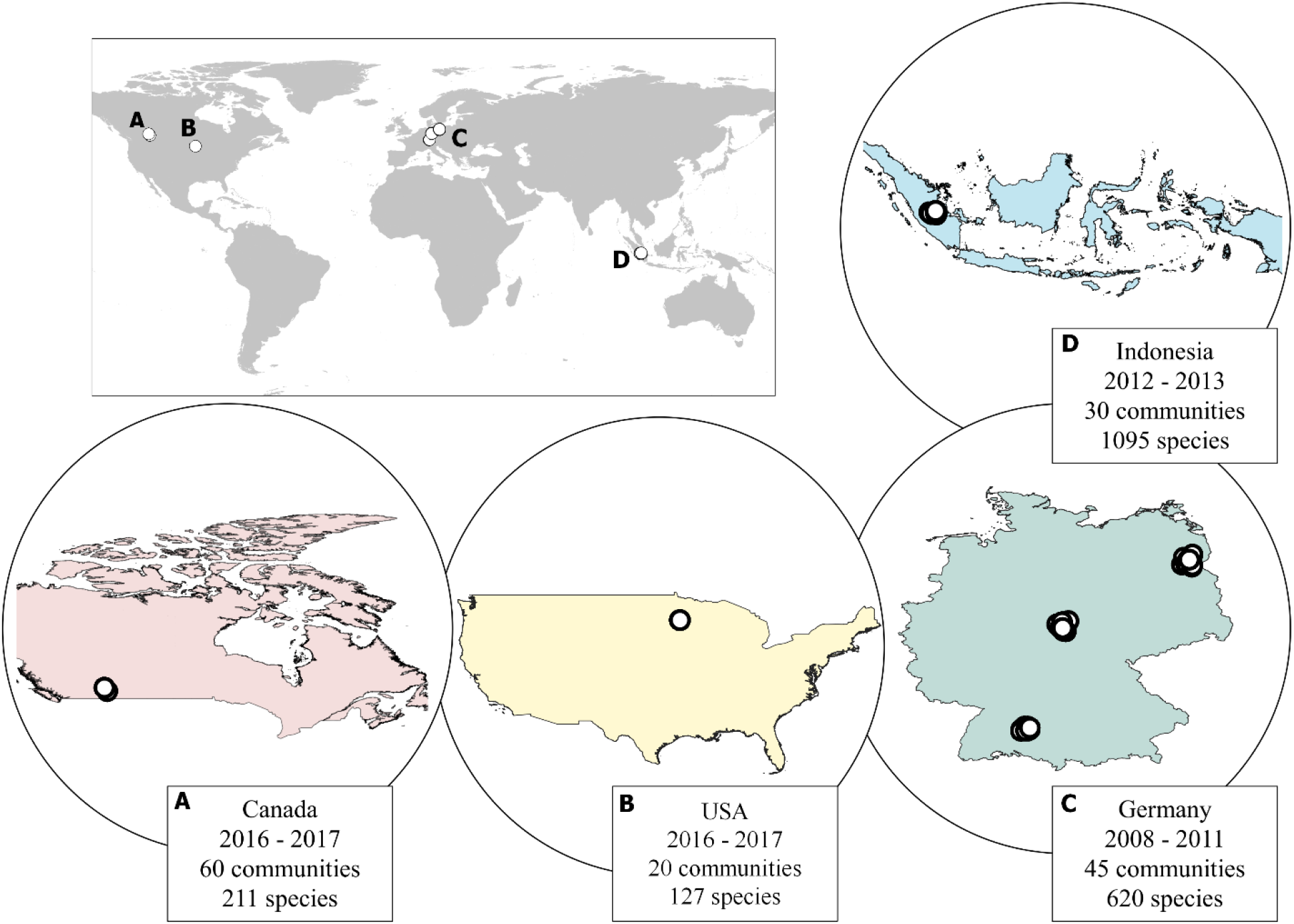
Distribution of study sites where 155 local soil communities were sampled to assess the site-specific body mass-abundance relationships.

### Sampling method

Soil samples were collected from litter and mineral soil layers from each study site. Standard sampling methods were applied in order to cover the meso- and macrofauna of the soil communities as comprehensively as possible: (1) sieving of leaf litter, hand sorting, large soil cores and heat extraction, and mustard extraction for sampling the macrofauna; (2) small soil cores and soil quadrats (16 × 16 cm, 5 cm depth) followed by heat extraction for the mesofauna. Invertebrates were either identified to the species level or classified as morphospecies. A detailed description of sampling methods at each site is provided in Table S1 (Supporting Information - S1). The different methods combined provide a comprehensive sample of the soil meso- and macrofauna community. Species abundances, population body masses, and the average body masses were calculated for each community (i.e. plot). Abundances were standardized and expressed as [individuals/m^2^]. For sites covering different sampling years, we averaged the abundances across sampling dates. Despite the three-dimensionality of the soil habitat, most of the animals are concentrated in the litter and top soil; therefore, we calculated the abundances in relation to the surface area, according to the conventionally used approach (Petersen and Luxton 1982, Ehnes et al. 2014).

### Environmental factors

We used the georeferences of the communities’ location and study year unit to extract soil annual mean temperature at a 1- km^2^ resolution for 0– 5 cm soil depth (Lembrechts et al. 2022). Additionally, other edaphic variables were used from the respective projects for each community: soil pH was measured using a digital pH meter, in CaCl_2_; water content in the soil was measured by comparing masses of dry and wet soil samples and expressed in % fresh weight; and carbon content was measured in the soil dry weight; Indonesia, USA and Canada: elemental analyser; Germany: automated CHNSO analyser. Data have been reported in detail in Ott et al. 2014, for Germany, Krashevska et al. 2015, for Indonesia, and Jochum et al. 2021b for USA and Canada sites.

### Statistical analyses

Prior to analyses, we excluded all larval or juvenile individuals from the data due to the complexity of identification to the species or genus level for juveniles. Subsequently, body mass and species-abundance data were log_10_-transformed to satisfy the assumptions of linearity of the analyses. After this log-transformation, the slope of their linear relationship equals the exponent of the power-law relationship in the untransformed data. In preparation of our further analyses, we independently ran linear regressions of the dependence of log_10_ abundance on log_10_ body mass for each of the 155 communities using the `lm` function in R (R Core Team, 2020). This resulted in a secondary dataset containing the 155 slopes of the local, within-community body mass-abundance relationships. Subsequently, we will refer to this data as the “slopes”. Additionally, we also calculated the log_10_ body-mass range for each community as the difference between the maximum and minimum body masses.

Thereafter, we used the ‘piecewise’ approach, based on confirmatory path analysis, to structural equation modeling (SEM) (Lefcheck 2016) to test the relative importance of the environmental variables and the body-mass range for the slopes of the local body mass-abundance relationships. This provides a mechanistic understanding of the direct and indirect pathways by which environmental conditions affect local body mass-abundance slopes and thus the distribution of abundances across small and large animals. We fitted the estimates within our SEM using Linear Mixed Effects Models, and we accounted for potential spatial autocorrelation by using the `corGaus` function from `nlme` package (Pinheiro et al. 2020), which required the use of a randomly parameterized dummy variable as a random effect (note that the `corGaus` function is only available for mixed-effects models that require a random effect variable). The initial model included the communities’ body-mass range, soil-carbon content and soil pH as direct effects on the slope, the soil temperature and soil water content as indirect effects, mediated by the local edaphic conditions and species body-mass range. While the SEM confirmed most of our prior hypotheses, it also identified an additional direct path from the soil temperature to the slope. The adequacy of this final model was determined by non-significant χ^2^ tests (P>0.05). All statistical analyses were performed using R Version 4.0.0 (R Core Team, 2020). We used `lme4` (version 3.1-150; Pinheiro et al. 2020) and the `piecewiseSEM` (version 2.1.0; Lefcheck, 2016) package to perform the Structural Equation Model.

As a sensitivity analysis, we also tested if other environmental characteristics have an impact on the abundance-mass slopes. In addition to the independent variables of our SEM analysis (soil temperature, soil water content, soil carbon content, soil pH and body-mass range), we also included additional independent variables (land-use intensity, human footprint index, litter layer mass and depth, C: N rate in the soil; see Supporting Information - S2 (Table S2) for detailed description of variables) in a linear mixed effects model (Supporting Information - S3, Table S3, Figure S3). This analysis indicated that none of the additional independent variables contributes to explaining variation in slopes.

To facilitate comparisons with prior studies, we also added an analysis of the general body mass-abundance relationship in a dataset pooling all local communities. This analysis shows the relative role of environmental and edaphic drivers (all independent variables as described in paragraph above) for species densities (dependent variable of the model instead of the slopes, Supporting information - S4, Table S4).

## Results

In the 155 local communities analyzed, body mass ranged from 0.000267 mg (*Liochthonius* sp. (Brachychthoniidae), Indonesia) to 6055 mg (*Lumbricus terrestris* (Lumbricidae), Germany) and species abundance ranged from 0.33 (*Uroballus koponeni* (Salticidae), *Carrhotus sannio* (Salticidae), Indonesia, among others) to 138,448 individuals/m^2^ (*Microppia minus* (Oppiidae), Germany). We found substantial variation in body mass-abundance slopes across the 155 local scaling relationships, ranging from -1.23 to -0.29 (Figure 2). Across all of the local communities’ slopes concentrated around the mean of -0.759 (Figure. 2, SD = 0.158, median = -0.770). Together, these findings indicate a stable global scaling relationship whose slope can be strongly modified locally.

**Figure 2.**
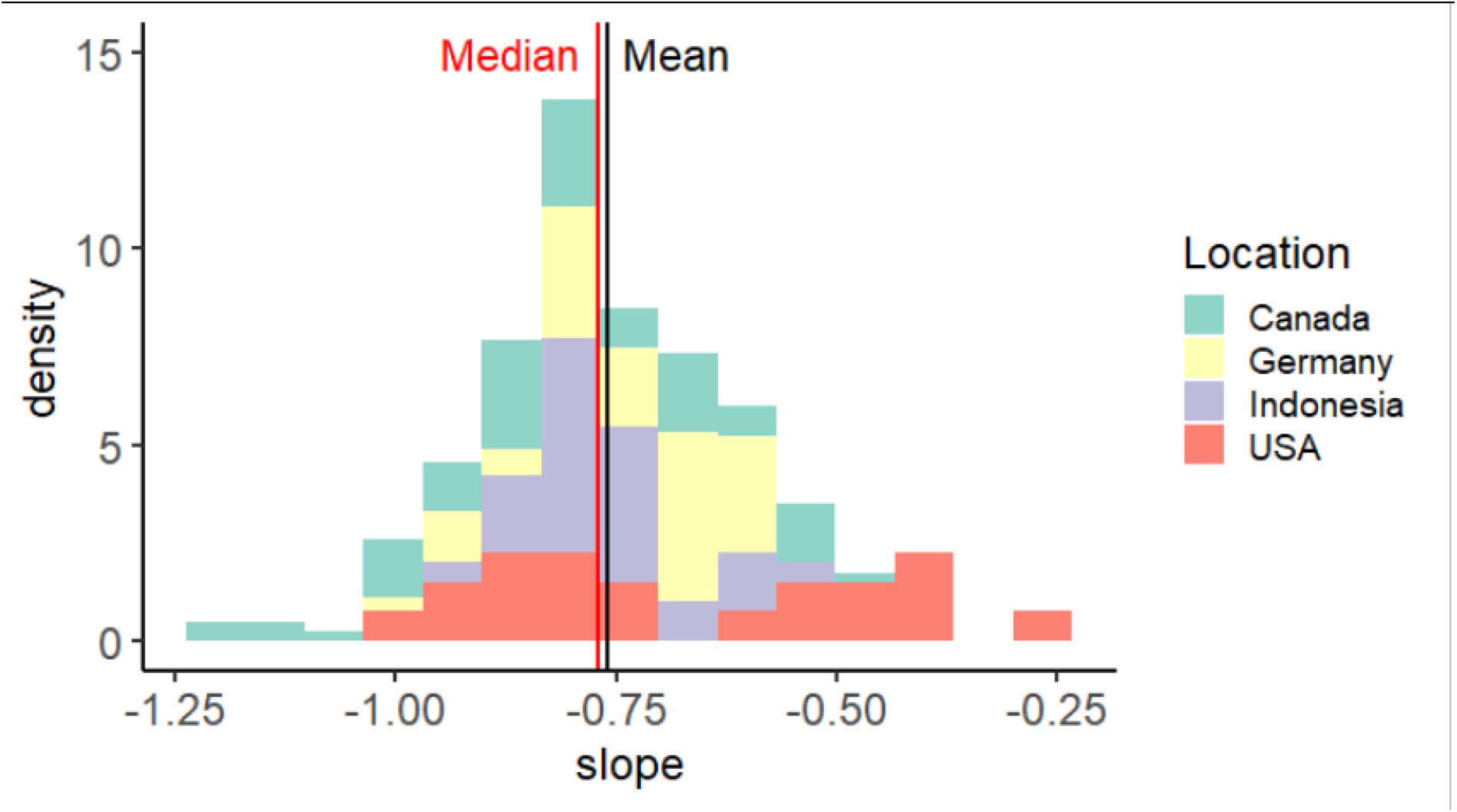
Frequency distribution of the slopes of the relationship between log_10_ body mass and log_10_ abundance for 155 local forest communities across three continents (mean = -0.759, SD = 0.158, median = -0.770).

Our SEM model adequately fit the data (Fisher’s C = 2.412; P-value = 0.661; effects of spatial autocorrelation have been accounted for in the model, Figure 3) and reveals direct positive effects of soil pH (path coefficient = 0.18), soil temperature (path coefficient = 0.21) and the body mass range (path coefficient = 0.52) as well as a direct negative effect of soil carbon content on the slope (path coefficient = -0.26). Positive and negative effects on the slope indicate shallower and steeper body mass-abundance-scaling relationships, respectively. Additionally, the SEM highlights important indirect effects of soil temperature (overall compound coefficient = 0.09) and water content (overall compound coefficient = - 0.07) on the slope. Soil water content increased the pH as well as the carbon content of the soil. Higher soil pH in turn had a positive effect on the slope. In contrast, higher carbon content decreased the slope value. Consequently, soil water content had an indirect positive effect on the slope mediated by pH (compound path coefficient = 0.03) and an indirect negative effect on the slope mediated by carbon content (compound path coefficient = -0.10). Soil temperature decreased soil pH and carbon content, as well as the body mass range. Hence, it has indirect negative effects on the slope via pH (compound path coefficient = -0.12) and the body mass range (compound path coefficient = -0.16), and an indirect positive effect via carbon content (compound path coefficient = 0.18). Additionally, the soil temperature has a direct positive effect on the slope (path coefficient = 0.21). Overall, our SEM analysis highlighted that the body mass range of the local communities has the strongest direct effect on the slope of the body mass-abundance relationship and reveals that soil temperature has much stronger indirect effects on the slope than soil water content.

**Figure 3.**
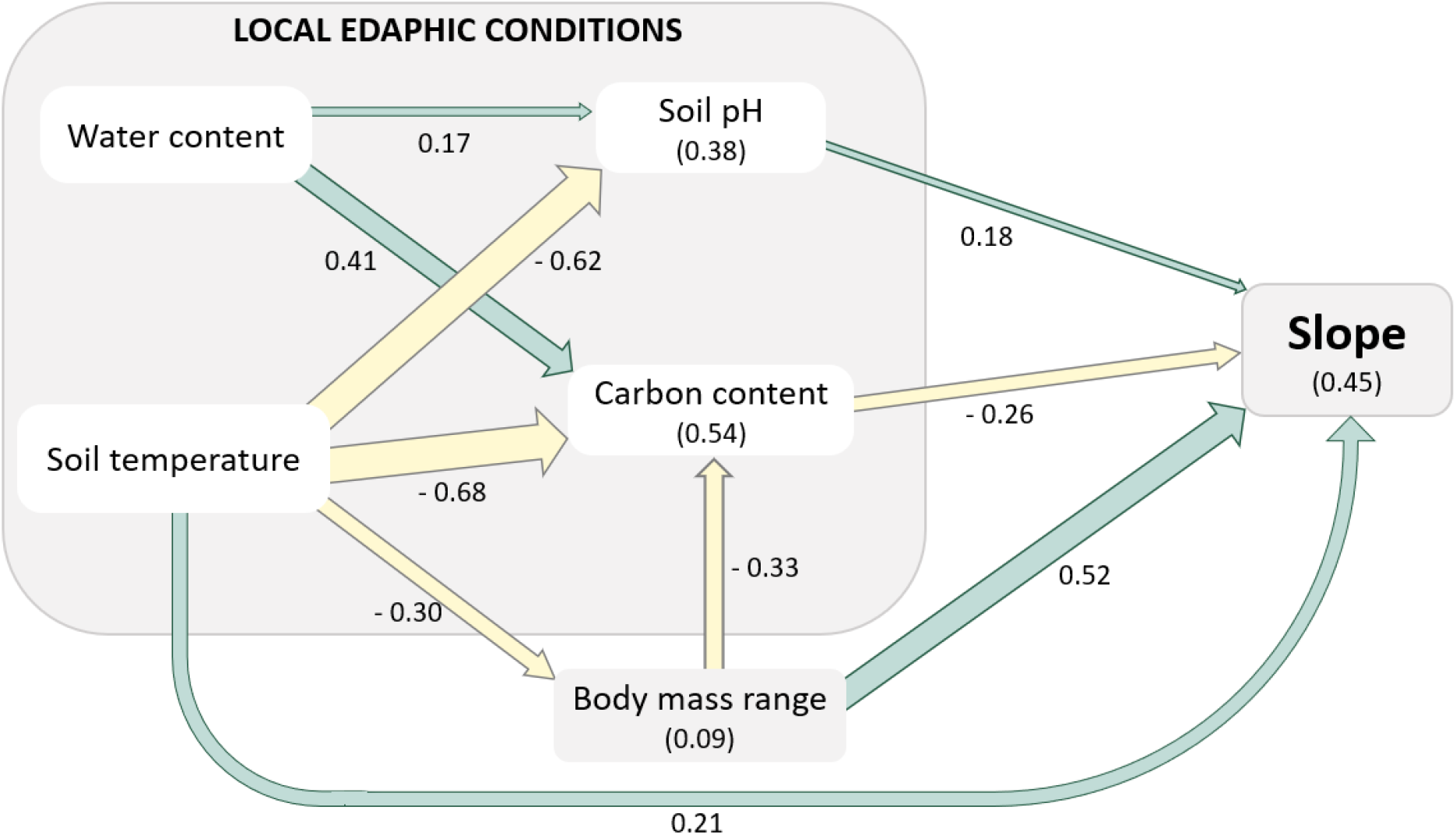
Structural equation model showing the direct and indirect effects of environmental variables on the local body mass-abundance slopes (Fisher’s C = 2.412; P-value = 0.661). Green and yellow arrows denote significant positive and negative effects respectively. Widths of arrows reflect standardized path coefficients (i.e. the relative strength of the individual effects) and indicate each predictor’s relative effect sizes. Numbers in parentheses inside the boxes indicate R-Squared values. Spatial autocorrelation effects accounted for by the model.

## Discussion

Our study disentangled the direct and indirect effects of environmental conditions on the body mass-abundance relationships of local soil-animal communities. Despite a global average abundance-mass scaling slope of -0.759, in line with theoretical expectations (Damuth 1981, 1987, White et al. 2007b) and prior empirical analyses of global relationships across communities (White et al. 2007b), we found substantial variation in these slopes across the 155 communities when analyzed separately (ranging between -1.23 and -0.29). Consistent with our hypothesis, we found that this variation in slopes can be explained by climatic conditions (soil water content and soil temperature) that exert strong direct and indirect effects via local edaphic (soil carbon content and soil pH) conditions and the body-mass range of the community. These results reinforce the importance of soil temperature and soil water content as strong environmental drivers of soil community structure (Phillips et al. 2019, van den Hoogen et al. 2019, Johnston and Sibly 2020), and illustrate how they influence the body mass-abundance structure of soil communities in concert with edaphic conditions. Our quantifications of these direct and indirect effects provide an important first step towards a mechanistic understanding of how soil communities respond to different environmental and edaphic conditions. These results also facilitate our understanding regarding future climatic scenarios, as shifts in the body-mass ranges and the altered distribution of abundances across size classes can have strong effects on population dynamics, community stability, and ecosystem functioning (Emmerson 2012, Brose et al. 2017a).

Our analysis revealed that the body mass range of the community is the strongest source of variation in the slopes of the body mass-abundance relationship of local communities (Figure 2). Research has shown that slope values vary widely when a narrow range of body mass is considered (White et al. 2007b, Hayward et al. 2010). However, these narrow body mass ranges are usually associated with smaller geographic scales or incomplete taxonomic samples, and may thus yield an artifactual component in the study’s results, mainly because it could indicate that the local communities are only partially included due to differences in sampling across sites. By contrast, our study comprised data of comprehensive belowground invertebrate communities that have all been sampled by the same combination of methods. Therefore, sampling biases are unlikely to be responsible for the variance in local body-mass ranges across communities and its effect on the body mass-abundance scaling slopes in our study. Instead, these ranges vary with shifts in the minima and maxima of the body masses in the communities (see Supporting Information S5, Figure S5).

Our study also showed that the edaphic conditions, soil pH and carbon content, have important direct effects on the slopes of the body mass-abundance relationships. Both edaphic factors have been shown to be important factors driving the general abundance of soil animals (Johnston and Sibly 2020). We show that a higher carbon content in the soil leads to steeper, more negative slopes, indicating relatively higher abundances of small compared to large animals. This matches research showing that soil systems with higher carbon contents are usually dominated by small soil animals, while the opposite is true for larger species (e.g. Chilopoda, Coleoptera, Clitellata) that occur in soils with lower carbon content (Johnston and Sibly 2020). Additionally, we found that increasing soil pH (i.e. soils becoming less acidic) leads to shallower slopes and thus benefits the large species in terms of abundance. Soil acidity is often associated with multiple nutrient availability (Binkley and Vitousek, 2000), and the species’ optimal pH ranges differ across phylogenetic groups. The soil macrofauna is usually restricted to soils with pH values above 3.5. Under the impact of acidification, soil-fauna individuals move downwards trying to mitigate the surface stress, altering the community composition and impacting ecosystem functions (e.g. organic-matter decomposition and greenhouse-gas emissions) (Wei et al. 2017). Catalase activity, the enzyme responsible for decomposing hydrogen peroxide into water and oxygen, decreases significantly with pH 6.5 to 4.0. The malfunction of this enzyme is lethal for organisms due to animal intoxication from the accumulation of H_2_O_2_, and injury of the cell structure membrane (Vitória et al. 2001). These findings of prior studies provide mechanistic explanations for our result that abundance-mass slopes are more steeply negative at the lower soil pH of the ecosystem. Overall, our study thus extends previous findings on the importance of pH and carbon content for soil animals by quantifying the distribution of abundance across different size classes.

We showed that soil temperature has a direct positive effect on the slope, which can be translated into a shift in relative densities from smaller to larger animals. The influence of soil temperature is directly related to species metabolism and resource requirements, which vary depending on the species size (Allen et al. 2002). An increase in temperature has a greater impact on smaller species, due to their relatively higher metabolic demands with increasing temperature, in comparison to larger species (Johnston and Sibly 2018, 2020). When experiencing higher metabolic demands, species are expected to increase resource uptake or, if this is not possible, exhibit declines in their population densities. Consistent with our results, Johnston and Sibly (2020) found that, under higher temperatures, smaller soil animal abundances declined, while, under low temperatures, larger soil animals experienced a decrease in their abundances. Our results extend this finding to within-community patterns and highlight that warming can cause a substantial reshuffling of abundance and thus biomass to the benefit of large species.

Furthermore, we show for the first time that the climatic variables soil temperature and water content exert important indirect effects on the local body mass-abundance scaling relationships. First, temperature leads to steeper slopes by decreasing the body mass range. While body masses generally decrease with warming, the maxima of body masses decrease more steeply than the minima, which is responsible for the decrease in range (Supporting information - S5, Figure S5). Second, temperature and water content both indirectly affect the body mass-abundance relationship of the communities by influencing soil pH and carbon content. Soil temperature is known to have a negative effect on the carbon content in the soil (Schimel et al. 1994), mainly by increasing soil carbon decomposition rates (Smith et al. 2008). Increasing soil temperature acts as the activation energy for the processes that effectively increase the carbon mineralization rate (Ågren and Wetterstedt 2007). In this context, rising temperatures will constrain the abundance of smaller species also by decreasing the carbon content in the soil. Overall, we showed that the dominant effects of both soil temperature and water content on the energy distribution across size classes within local communities are indirectly mediated via changes in edaphic conditions and the body-mass range realized in the community.

## Conclusions

Our study confirmed a three-quarter power-law scaling of population abundance with body mass when averaged across soil animal communities of four locations of the globe. However, we also found substantial variation in the power-law exponents along environmental gradients. Specifically, we addressed the consequences of variation in soil temperature and water content. Our study showed a net positive effect of soil temperature on the slope of the body mass-abundance relationship, which is mainly due to the combination of the direct positive effect with the indirect positive effect via soil carbon content and the indirect negative effect via body mass range. This implies that warming generates a less negative slope, resulting in a relative redistribution of biomass from small to large species. Furthermore, the negative indirect effect of soil water content via soil carbon content is roughly three times stronger than the positive indirect effect via soil pH, which yields a negative net effect on the body mass-abundance slopes. This implies that increasing soil water content yields steeper, more negative slopes and thus favors small over large species in the communities. Together, these results reveal the important indirect constraints of soil-climatic variables on the distribution of abundances across size classes, which explains the substantial variation in local body mass-abundance relationships.

Future climate projections suggest a scenario of decreasing precipitation rates and increasing temperatures for most global regions (IPCC 2022, In Press). Therefore, lower contents of water in the soils and increased soil temperatures are expected. In combination with our results, this implies that belowground communities will experience a shift in the biomass distribution from smaller to larger species, a reflex of the shallower slopes observed in this study. While our study also corroborates an overall decrease in average body mass with warming (Daufresne et al. 2009), our results suggest that abundance, and thus energy, is shifting from the smaller to the largest species in the community. Together, these findings imply that warming benefits the small when analyzed across communities (i.e. the shift to lower body masses) but it also benefits the larger species within communities (shifts in biomass to the larger species of the community). Such increasing dominance of large species in the community has several implications, including (1) dominance of slow energy channels, especially when large decomposers are advantaged, and (2) increasing top-down control as densities of large predators increase. The slower energy channels, composed of large species, are characterized by slower population dynamics, longer material retention times and, in many cases, fungal resources at the base (Moore et al. 2003). Increasing densities of large predators with their high per capita feeding rate can be indicative of increasing feeding rates (Rall et al. 2012) (Schneider et al. 2012), which yields higher interaction strength and energy fluxes through the food webs. Both a shift in the balance between energy channels and increased top-down control have the potential to destabilize community dynamics (Johnson et al. 2014, Jacquet et al. 2016, Wolkovich 2016, Brose et al. 2017b, Zhou et al. 2022). However, this could be offset by the generally stabilizing effect of large species on food-web dynamics (Brose et al. 2006, 2017b, Heckmann et al. 2012). Similarly, increases in the biomass of large species may promote ecosystem functioning at the base of the food web if maximum trophic levels and omnivore rates are increased (Schneider et al. 2012, Wang and Brose 2018, Wang et al. 2019). Interestingly, this suggests that integrating our findings on biomass distribution shifts with food-web approaches offers great potential for predicting the community-level consequences of future warming and drought.

Overall, our study revealed the complex interplay between soil temperature and soil water content and their effects on the body mass-abundance structure of soil communities, which facilitates future modeling approaches to predict the consequences of global change for soil communities and their functioning. Together, this will be an important step towards a mechanistic and predictive understanding of how soil community dynamics and functioning are expected to respond to global change.

## Data archiving statement

Data supporting the results will be available from the Dryad Digital Repository

