## Supporting Information for "Global environmental drivers of local abundance-mass scaling in soil animal communities"

**Supporting Information – S1**

*Table S1: description of sampling methods in each site*

| Site Location | Group | Sampling method | Number of plots | Area per plot m2 | Soil core | Sampling date | Body mass estimation |
|---|---|---|---|---|---|---|---|
| USA, Canada | Macrofauna | litter sieving, hand sorting | 80 | 0.5 | NA | 2016 - 2017 | length-mass regressions from Wardhaugh 2013, Sohlström et al. 2018 |
| USA, Canada | Mesofauna | 1 soil core, heat-extraction | 80 | 0.00196 | 5cm diameter, 10 cm depth | 2016 - 2017 | length–mass regressions for specific taxa from Mercer et al. 2001 |
| Germany | Macrofauna | 2 soil cores, heat-extraction | 48 | | 20cm diameter, two samples per plot | 2008 - 2011 | measured or estimated with mass-length regressions from Ehnes et al. 2014 |
| Germany | Macrofauna | litter sieving, mustard extraction | 48 | 0.25 | NA | 2008 - 2011 | measured or estimated with mass-length regressions from Ehnes et al. 2014 |
| Germany | Mesofauna | 2 soil cores, heat-extraction | 48 | | 5cm diameter, two samples per plot | 2008 - 2011 | measured or estimated with mass-length regressions from Ehnes et al. 2014 |
| Indonesia | Mesofauna | 2 soil cores, heat-extraction | 32 | 0.0256 | litter + 5cm depth | 2013 | length-mass regressions were used for Collembola: dry weight (Peterson 1975) |
| Indonesia | Macrofauna | litter sieving, heat-extraction | 32 | 3 | NA | 2012 | length-body mass regressions were used to estimate spp body mass (Sohlström et al. 2018) |

**Supporting Information – S2**

*Additional environmental variables descriptors*

In order to explore the effect of additional environmental and edaphic variables on our analysis, we extracted the human footprint index based on data on human pressures at 1 $km^2$ resolution (from 1993 and 2009) (Venter et al. 2016). Current global scale land-change classifications were extracted from van Asselen and Verburg (2012) at a 5-arcminute resolution (Table S2*)*. Original land-use maps were converted to numerical data, following Pouzols et al. (2014) and Eitelberg (2018), with values imputation for the missing categories (Table 1 - S2). Other environmental variables were available from the respective projects for each community: litter layer was measured (cm) and weighted (g/m2); carbon and nitrogen content were measured in the soil (dry weight), and used to calculate C: N ratio. We used the georeferences of the communities' location and study years in a 0.05 degrees unit to extract NDVI (from 2000 to 2018) (MOD13C2 Series – Didan, 2015).

*Table S2: Current global scale land-change classifications were extracted from Van Asselen & Verburg (2012) and Eitelberg (2018)*

| Land System | Pouzols et al. (2014) | Eitelberg (2018) | Imputation | Final intensity value |
|---|---|---|---|---|
| Cropland; extensive, few livestock | 0.4 | 0.4 | | 0.4 |
| Cropland; extensive, bovines, goats & sheep | | 0.4 | | 0.4 |
| Cropland; extensive, pigs & poultry | | | 0.45 | 0.45 |
| Cropland; medium intensive, few livestock | 0.3 | 0.3 | | 0.3 |
| Cropland; medium intensive, bovines, goats & sheep | | 0.3 | | 0.3 |
| Cropland; medium intensive, pigs & poultry | | | 0.35 | 0.35 |
| Cropland; intensive, few livestock | 0.2 | 0.2 | | 0.2 |
| Cropland; intensive, bovines, goats & sheep | | 0.2 | | 0.2 |
| Cropland; intensive, pigs & poultry | | | 0.25 | 0.25 |
| Mosaic cropland and grassland; bovines, goats & sheep | | 0.8 | | 0.8 |
| Mosaic cropland and grassland; pigs & poultry | | | 0.85 | 0.85 |
| Mosaic cropland (ext.) and grassland; few livestock | 0.7 | 0.7 | | 0.7 |
| Mosaic cropland (med. int.) and grassland; few livestock | 0.6 | 0.6 | | 0.6 |
| Mosaic cropland (int.) and grassland; few livestock | 0.5 | 0.5 | | 0.5 |
| Mosaic cropland and forest; pigs & poultry | | | 0.55 | 0.55 |
| Mosaic cropland (ext.) and forest; few livestock | 0.7 | 0.7 | | 0.7 |
| Mosaic cropland (med. int.) and forest; few livestock | 0.6 | 0.6 | | 0.6 |
| Mosaic cropland (int.) and forest; few livestock | 0.5 | 0.5 | | 0.5 |
| Dense forest | 1 | 1 | | 1 |
| Open forest, few livestock | 0.9 | 0.9 | | 0.9 |
| Open forest, pigs & poultry | | | 0.95 | 0.95 |
| Mosaic grassland and forest | 1 | 1 | | 1 |
| Mosaic grassland and bare | 1 | 1 | | 1 |
| Grassland, natural | 1 | 1 | | 1 |
| Grassland, few livestock | 0.9 | 0.9 | | 0.9 |
| Grassland, bovines, goats & sheep | | 0.9 | | 0.9 |

| | | | |
|---|---|---|---|
| Bare | 0.1 | 1 | 1 |
| Bare, few livestock | 0.9 | 0.9 | 0.9 |
| Peri-urban & villages | 0.1 | 0.1 | 0.1 |
| Urban | 0.1 | 0 | 0 |

**Supporting Information – S3**

To evaluate if additional environmental variables affect body mass-abundance relationships across local communities, we used Linear Mixed Effects Models that relate the previously evaluated slopes of the body mass-abundance relationship for each soil animal community to the local community's body mass range and environmental variables (soil temperature, precipitation, land-use intensity, soil pH, human footprint index, the carbon content in the soil, litter layer mass and depth, C: N rate in the soil, water content in the soil). Based on a correlation analysis of all environmental variables, we removed NDVI from the model due to its high correlation with soil temperature. The mixed-modeling approach was used to account for potential spatial autocorrelation by using the `corGaus` function from `nlme` package (Pinheiro et al. 2020), which required the use of a randomly parameterized dummy variable as a random effect (note that the `corGaus` function is only available for mixed-effects models that require a random effect variable). Each of the independent variables was added as a linear term, without interactions. We started with the full model comprising all independent variables and selected the best-fitting model by the 'dredge' function of the `MuMIn` package (Barton 2022), using the Bayesian information criterion (BIC) for model comparison ($\Delta$BIC <2).

The two most supported models ($\Delta$BIC <2) were used to generate model-averaged estimates of the parameters using the 'model.avg' function from the `MuMIn` package. Model-averaged estimates from the top models ($\Delta$BIC <2) included the body-mass range, water content in the soil, soil carbon content and temperature. This final model reveals linear increases in the slope with increasing body-mass range, soil temperature and water content and decreases with increasing soil carbon content (Table 1 - S3). The general relationships between the slopes and the variables selected in the final model were illustrated in Figure 1 (S3).

*Table S3: Summary of the parameter estimates of the final Mixed-Effect Model (conditional average) for slope prediction. Estimates, standard errors and p-value for the Z-statistic are indicated.*

| **Predictors** | **Estimates** | **Std. Error** | **Pr(>\|z\|)** |
|---|---|---|---|
| (Intercept) | -0.63931 | 0.15688 | $4.72 \times 10^{-5}$ |
| log body mass range | 0.10926 | 0.01452 | $< 2 \times 10^{-16}$ |
| log carbon content | -0.16443 | 0.03951 | $3.61 \times 10^{-5}$ |
| log soil temperature | 0.13989 | 0.05617 | 0.0135 |
| soil pH | 0.03220 | 0.01321 | 0.0156 |

*Figure S3: Relationships between the slopes of the body mass-abundance relationship in the communities in each location (colored symbols) with* ***A.*** *mean soil temperature* ($log_{10}$)*,(y = -0.81 + 0.051x,* $R^2$ *= 0.0063),* ***B.*** *soil pH, (y = -0.79 + 0.0066x,* $R^2$ *= 0.0015),* ***C.*** *body mass range of the communities* ($log_{10}$)*, (y = -0.68 + 0.13x,* $R^2$ *= 0.35),* ***D.*** *soil carbon content* ($log_{10}$)*, (y = -0.56 - 0.24x,* $R^2$ *= 0.18) and* ***E.*** *water content in the soil (% fresh weight), (y = -0.71 – 0.0017x,* $R^2$ *= 0.0014).*

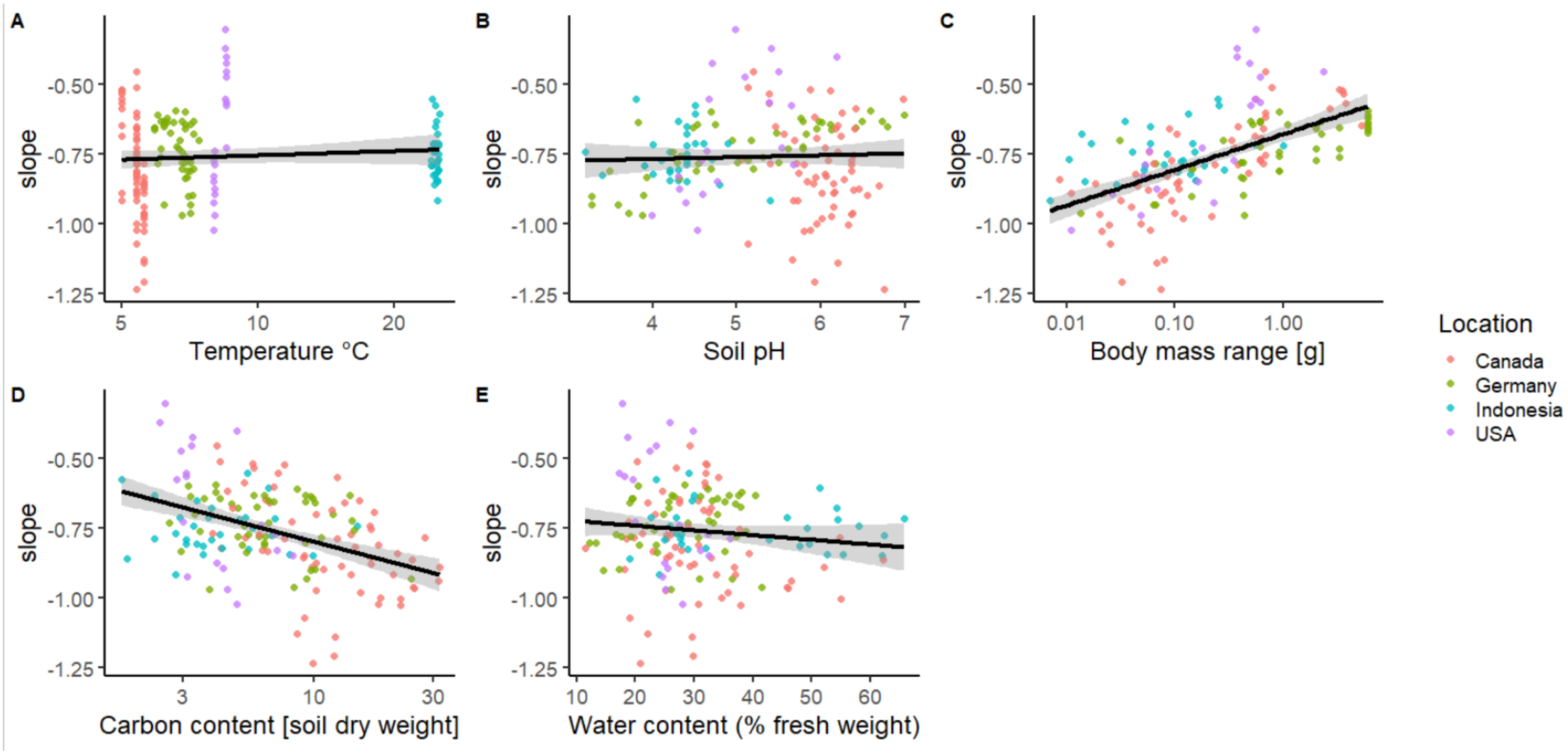

**Supporting Information – S4**

To describe general body mass-abundance relationships across communities, we fitted a linear model pooling the abundance and mass data of the species for all sites. We ran a linear regression of the dependence of each species $\log_{10}$ abundance on the $\log_{10}$ body mass and edaphic variables (soil temperature, precipitation, land-use intensity, soil pH, human footprint index, the carbon content in the soil, litter layer mass and depth, C: N rate in the soil, water content in the soil). Based on a correlation analysis of all environmental variables, we removed NDVI from the model due to its high correlation with soil temperature. Each of the independent variables was added as a linear term, without interactions. We started with the full model comprising all independent variables and selected the best-fitting model by the 'dredge' function of the `MuMIn` package (Barton 2022), using the Bayesian information criterion (BIC) for model comparison ($\Delta$BIC <2).

The two most supported models ($\Delta$BIC <2) were used to generate model-averaged estimates of the parameters using the 'model.avg' function from the `MuMIn` package. Model-averaged estimates from the top models ($\Delta$BIC <2) included body mass, human footprint index, land-use intensity, litter layer depth, soil temperature, soil pH and water content in the soil). This final model reveals linear increases in the species abundance with increasing human footprint index and land-use intensity and decreases with increasing species body mass, litter layer depth, soil temperature, soil pH and water content in the soil (Table 2 - S2). Our model can be used in future predictions to assess the abundance of soil species for large-scale projections.

*Table S4: Summary of the parameter estimates of the final Mixed-Effect Model for species abundances prediction. Estimates, standard errors and p-value for the Z-statistic are indicated.*

| **Predictors** | **Estimates** | **Std Error** | **Pr(>\|z\|)** |
|---|---|---|---|
| (Intercept) | 1.882579 | 0.124495 | $< 2 \times 10^{-16}$ |
| human footprint index | 0.006876 | 0.001473 | $3.07 \times 10^{-6}$ |
| land-use intensity | 0.246828 | 0.038845 | $< 2 \times 10^{-16}$ |
| litter layer depth | -0.020904 | 0.006052 | 0.000553 |
| log soil temperature | -2.818322 | 0.050278 | $< 2 \times 10^{-16}$ |
| log body mass | -0.743859 | 0.006296 | $< 2 \times 10^{-16}$ |
| soil pH | -0.090023 | 0.011250 | $< 2 \times 10^{-16}$ |
| log water content | -0.162260 | 0.058978 | 0.005945 |

**Supporting Information – S5**

To evaluate how the body-mass range of the communities varies along the gradient of temperature, we ran linear regressions of the dependence of **A.** $\log_{10}$ minimum body mass (g), **B.** $\log_{10}$ maximum body mass (g) and **C.** $\log_{10}$ body-mass range (g) (difference between maximum and minimum body masses) on soil temperature (°C) for each of the 155 communities using the `lm` function in R (R Core Team, 2020).

*Figure S5: Relationships between the soil temperature (log$_{10}$) in each location (colored symbols) with* ***A****. minimum body mass (log$_{10}$) (y = -4.4 – 1.3x, $R^2$ = 0.63)* ***B.*** *maximum body mass (log$_{10}$) (y = 0.12 – 0.83x, $R^2$ = 0.078) and* ***C.*** *body mass range (log$_{10}$) (y = 0.12 – 0.83x, $R^2$ = 0.078) in each community.*

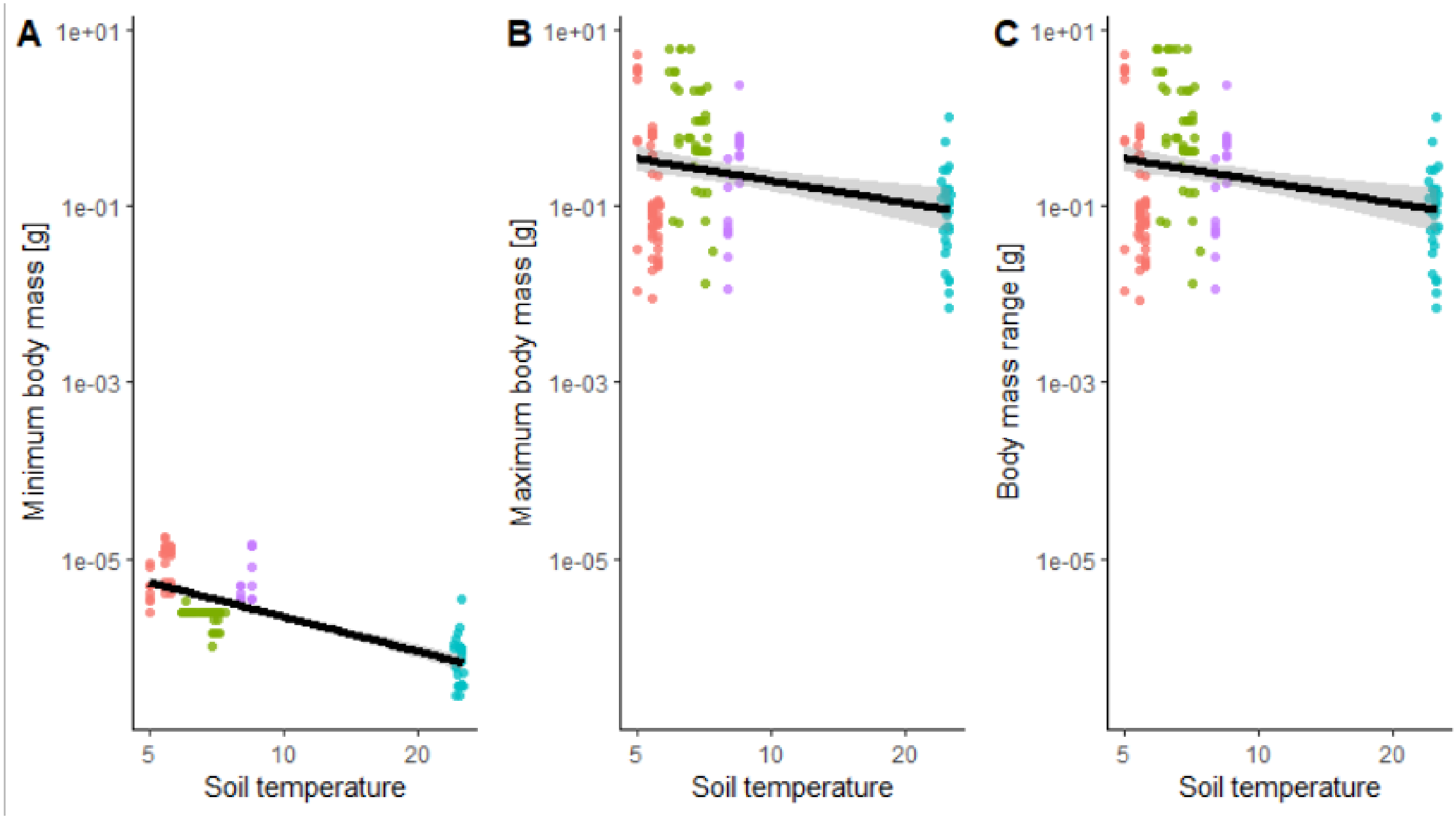
